# Identification of small molecule antivirals against HTLV-1 by targeting the hDLG1-Tax-1 protein-protein interaction

**DOI:** 10.1101/2023.03.15.532789

**Authors:** Sibusiso B. Maseko, Yasmine Brammerloo, Inge Van Molle, Adrià Sogues, Charlotte Martin, Christoph Gorgulla, Julien Olivet, Jeremy Blavier, Thandokuhle Ntombela, Frank Delvigne, Haribabu Arthanari, Kourosh Salehi-Ashtiani, Han Remaut, Steven Ballet, Alexander N. Volkov, Jean-Claude Twizere

## Abstract

Human T-cell leukemia virus type-1 (HTLV-1) is the first pathogenic retrovirus discovered in human. Although HTLV-1-induced diseases are well characterized and linked to the encoded Tax-1 oncoprotein, there is currently no strategy to target Tax-1 functions with small molecules. Here, we analysed the binding of Tax-1 to the human homolog of the drosophila discs large tumor suppressor (hDLG1/SAP97), a multi-domain scaffolding protein involved in Tax-1-transformation ability. We have solved the structures of the PDZ binding motif (PBM) of Tax-1 in complex with the PDZ1 and PDZ2 domains of hDLG1 and assessed the binding of 10 million molecules by virtual screening. Among the 19 experimentally confirmed compounds, one systematically inhibited the Tax-1-hDLG1 interaction in different biophysical and cellular assays, as well as HTLV-1 cell-to-cell transmission in a T-cell model. Thus, our work demonstrates that interactions involving Tax-1 PDZ-domains are amenable to small-molecule inhibition, which provides a framework for the design of targeted therapies for HTLV-1-induced diseases.

## 1. Introduction

Human T-cell lymphotropic viruses (HTLV-1 to −4) are members of the *Deltaretrovirus* genus of the *Retroviridae* family. The first human retrovirus to be isolated 42 years ago (Poiesz et al., 1981), HTLV-1 is the etiological agent of adult T-cell leukemia/lymphoma (ATL), an aggressive neoplasm, as well as HTLV-1-associated myelopathy (HAM). Also termed tropical spastic paraparesis (TSP), HAM is a degenerative neurologic syndrome. Currently, about 10-20 million people worldwide are infected with HTLV-1, and 1 to 10% of them will develop severe ATL or HAM/TSP diseases (Gessain et al., 1985; Hinuma et al., 1981; Pasquier et al., 2018). For the last years, ATL patients have been treated using chemotherapy-based approaches, but with very limited benefit as the median survival rate is only 8-10 months (Bazarbachi et al., 2011; Cook and Phillips, 2021; Utsunomiya et al., 2015). Improved survival is achieved by antiviral therapies combining zidovudine and interferon-alpha (Nasr et al., 2017), allogenic hematopoietic stem cell transplantation (Fuji et al., 2016), or the use of monoclonal antibodies targeting CC chemokine receptor 4 (CCR4), which is frequently expressed in ATL patient samples (Ishida et al., 2015; Yoshie et al., 2002). However, the above therapies are not fully effective, mainly due to clinical disease heterogeneity, discrepancies in treatment between countries, and lack of specific and universally targeted drugs (Cook and Phillips, 2021). In contrast to ATL, no treatment is available for HAM/TSP patients. Both HTLV-1-induced diseases are associated with high proviral loads. Targeting viral replication, including the re-positioning of existing anti-human immunodeficiency virus (HIV) drugs, was investigated to decrease HTLV-1 cell-to-cell transmission (Pasquier et al., 2018) and may lead to effective anti-HTLV-1 therapies.

In addition to the essential retroviral genes, HTLV-1 encodes the *Tax* and *HBZ* viral oncogenes which interfere with transcription and post-transcription regulation towards pathogenesis, and are able to induce leukemia-lymphoma in transgenic mice (Hasegawa et al., 2006; Ohsugi et al., 2007). However, there is a difference in expression kinetics of sense transcription (e.g *Tax* expression), which is heterogeneous and intermittent, and antisense transcription (e.g., *HBZ* expression), which is more stable (Mohanty and Harhaj, 2020). The Tax-1 protein is indeed a potent transcriptional activator of viral and cellular genes through association with transcription modulators (Giam and Semmes, 2016) and a major determinant in HTLV pathogenesis and persistence (Mohanty and Harhaj, 2020).

Tax-1 is a multi-domain, modular protein reported to interact with more than 200 cellular protein targets (Vandermeulen et al., 2021). The C-terminus of Tax-1 harbours a PDZ-binding motif (PBM), which confers binding to a class of proteins containing a defined structure of ~90 amino acids known as PDZ (PSD-95/Discs Large/ZO-1) domain (Harris and Lim, 2001; Tonikian et al., 2008). PDZ-containing proteins are modular polypeptides implicated in the assembly of large protein complexes, mediating signalling, cell polarity, and communication (Hung and Sheng, 2002). Our analysis of the human genome estimates the presence of 256 PDZ domains in 149 distinct proteins, excluding variants and isoforms (Belotti et al., 2013). To date, 14 PDZ proteins interacting with HTLV-1 Tax have been identified (Boxus et al., 2008; Yan et al., 2009), and may have important implications in HTLV-1-induced leukemogenesis process. In particular, it was shown that the Tax-1 PBM motif, which is not present in the HTLV-2 Tax counterpart, promotes transformation of rat fibroblasts and mouse lymphocytes *in vitro* (Higuchi et al., 2007; Hirata et al., 2004; Tsubata et al., 2005), as well as persistence of the HTLV-1 virus *in vivo*(Pérès et al., 2018).

In this study, we hypothesized that disrupting Tax-1-PDZ interactions would provide novel anti-HTLV-1 pathogenesis strategies, which could potentially be used to combat ATL and TSP/HAM diseases. As a proof-of-concept, we targeted the binding of Tax-1 PBM to the human homolog of the drosophila discs large tumor suppressor (hDLG1/SAP97), the first isolated PDZ-containing binding partner of Tax-1. Using a range of structural biology techniques, we characterized Tax-1 PBM interactions with hDLG1 PDZ domains, performed virtual screening of ultra-large libraries of small chemical compounds, and validated the hits in biophysical and cellular assays. One of the identified small molecules can disrupt Tax-1/hDLG1 interaction and inhibits HTLV-1 cell-to-cell transmission. This work showcases an attractive antiviral drug discovery pipeline, which should be readily applicable to target other protein-protein virus-host interactions.

## 2. Materials and Methods

### 2.1. GST-pulldowns

GST-fused hDLG1 PDZ domains were expressed in *E. coli*, while Tax-1 was obtained from culturing HTLV-1 infected cells (MT2). The PDZ domains were then purified using glutathione Sepharose beads. The beads, containing the fusion proteins, were then incubated overnight with Tax-1 containing lysates. The beads were washed next and subjected for western blot analysis with an α-Tax-1 antibody.

### 2.2. Gaussia princeps luciferase complementation (GPCA) and mammalian nanoluciferase-2 hybrid (mN2H) assays

HEK293 cells were seeded in 24-well plates, then transfected in triplicates with GL1/GL2 (GPCA) or N1/N2 (mN2H) plasmids expressing the fusion proteins 24 hours later. (Nano)luciferase activity was measured on cells or lysates in 96-well plates for mN2H and GPCA respectively. The results were normalized relative to the value of the (nano)luciferase control. The normalized luciferase ratio (NLR) was calculated as follows: NLR = co-transfection luciferase value (GL1 + GL2) / (GL1 luciferase value alone + GL2 luciferase value alone). An interaction is considered positive or validated when NLR ≥ 3.5. For small molecule testing, the compounds in different concentrations were added 24-hour post transfection. Read outs were done as above. Cell viability was measured using Cell Titer Glo kit from Promega (G9681). Reduction in luciferase counts due to the small molecule was normalised to the cell viability.

### 2.3. Synthetic Peptides

A biotinylated derivative of the (10 amino acid-long) C-terminus of Tax-1 (i.e. Biotin-Ahx-SEKHFRETEV-OH) was prepared by Fmoc-based solid-phase peptide synthesis and purified by high-performance liquid chromatography to >99% purity (see Supplementary Text 1 for detailed synthesis protocol and characterization). All other Tax-1-derived peptides were synthesized by Biomatik, Canada, as acetate salts of >98% purity.

### 2.4. Protein expression and purification

Plasmids harbouring hDLG1 PDZ1, PDZ2 or PDZ1+2 tandem were obtained from Addgene (Tonikian et al., 2008) or purchased from GeneScript respectively and transformed into *E. coli* BL21. For expression, the culture was grown at 37°C in LB, shaking (180 rpm) until OD_600_ reached 0.4-0.6 then induced by addition of 1.0 mM IPTG and further grown for 4 hrs. The isotopically labelled U-[^13^C,^15^N]or U-[^15^N] proteins were produced in minimal medium following a published protocol (Volkov et al., 2013). GST-fused proteins were purified using glutathione Sepharose 4B beads (Cytiva). Following GST tag removal, the proteins were further purified using a HiPrep 16/60 Sephacryl S-100 HR size exclusion column.

### 2.5. Isothermal titration calorimetry (ITC)

Measurements were carried out on an Microcal ITC200 calorimeter at 25 °C in 10 mM Tris pH 8.0. The sample cell contained 20 μM protein and syringe had 250 μM peptide. Titrations comprised 26 × 1.5 μL injections of peptide into the protein, with 90 s intervals. An initial injection of 0.5 μL was made and discarded during data analysis. The data were fitted to a single binding site model using the Microcal LLC ITC200 Origin software.

### 2.6. Nuclear magnetic resonance (NMR) Spectroscopy

All NMR spectra were acquired at 298 K in 20 mM sodium phosphate 50 mM NaCl pH 6.0, 2 mM DTT and 10 % D_2_O for the lock on a Bruker Avance III HD 800 MHz spectrometer (equipped with a TCI cryoprobe) or a Varian Direct-Drive 600 MHz spectrometer (fitted with a PFG-Z cold probe). The data were processed in NMRPipe (Delaglio et al., 1995) and analyzed in CCPNMR (Vranken et al., 2005). The assignments of backbone resonances of the individual hDLG1 PDZ1 and PDZ2 domains were obtained from 0.7-1.1 mM U-[^13^C,^15^N] protein samples using a standard set of 3D BEST HNCACB, HN(CO)CACB, HNCO and HN(CA)CO experiments and aided by the published assignments (Cierpicki et al., 2005; Liu et al., 2007; Tully et al., 2012). Subsequently, the NMR assignments of the individual PDZ domains were transferred to the spectra of hDLG1 PDZ1+2 tandem and verified by standard triple-resonance experiments. The NMR chemical shift perturbation experiments were performed by incremental addition of 10 mM stock solutions of Tax-1 peptides to U-[^15^N] or U-[^13^C,^15^N] protein samples at the initial concentration of 0.3 mM. At each increment, changes in chemical shifts of the protein resonances were monitored in 2D [^1^H,^15^N] HSQC spectra. The average amide chemical shift perturbations (Δδ_avg_) were calculated as Δδ_avg_ = (Δδ_N_^2^/50+Δδ_H_^2^/2)^0.5^, where Δδ_N_ and Δδ_H_ are the chemical shift perturbations of the amide nitrogen and proton, respectively.

### 2.7. X-ray crystallography

Crystallization screens were set up with purified hDLG1 PDZ1 (10.6 mg/mL) or PDZ2 (11.5 mg/ml) and 4 molar equivalents of Tax-1 ETEV peptide using the sitting drop vapour diffusion method and a Mosquito nanolitre-dispensing robot at room temperature (TTP Labtech, Melbourn, UK). The best hDLG1 PDZ1-ETEV crystals appeared after 3-5 days in 0.1 M Bis-Tris pH 5.5, 0.18 M (NH_4_)2SO_4_, 24% PEG3350. Similarly, diffraction-quality crystals of hDLG1 PDZ2-ETEV complex appeared after 7-10 days in 0.1 M Bis-Tris pH 5.5, 0.075 M (NH_4_)2SO_4_, 24% PEG3350. The crystallization buffer was supplemented with 10% glycerol, and crystals were mounted in nylon loops and flash-cooled in liquid nitrogen.

X-ray diffraction data were collected at 100 K using the Beamline Proxima 2 at the Soleil synchrotron (Gif-sur-Yvette, France) and the Beamline iO4 at Diamond Light Source (Didcot, UK). The diffraction data were processed with AutoProc (Clemens et al., 2011). The crystals of hDLG1 PDZ2-ETEV presented high levels of anisotropy, necessitating further data processing with Staraniso (Global Phasing suite). The crystal structures were determined by molecular replacement with phaser (Bunkóczi et al., 2013; Liebschner et al., 2019) and PDB 3RL7 as the search model. The structures were refined through iterative cycles of manual model building with COOT (Emsley and Cowtan, 2004) reciprocal space refinement with phenix.refine (Afonine et al., 2012) and Buster (Smart et al., 2012). The crystallographic statistics are shown in Supplementary Table 1. Atomic coordinates and structure factors of hDLG1 PDZ1 and PDZ2 complexes with ETEV peptide have been deposited in the protein data bank (PDB) under the accession codes 8CN1 and 8CN3, respectively.

### 2.8. Virtual screening

The virtual screening was carried out with VirtualFlow (Gorgulla et al., 2020). The docking was performed with QuickVina 2 (Alhossary et al., 2015), with the exhaustiveness set to 4. The docking box had a size of 15.0 x 18.0 x 15.0 Å, and the protein was held rigid. The protein was prepared with Schrödinger Maestro and AutoDockTools (Maestro, 2020; Morris et al., 2009). The computations were carried out in the Google Cloud (Morris et al., 2009). After the screening, the best hits were post processed with DataWarrior, and the compounds with a logP > 5 and a M > 600 Da were removed from the docking score list.

### 2.9. Fluorescence polarization (FP)

All experiments were performed with GST-hDLG1 PDZ1 or PDZ2 proteins in NMR buffer in 384 well-plates. For the compound screening, the samples contained 3.0 μM GST-hDLG1 PDZ1, 1 mM compound (added from 20 mM stock in DMSO) and incubated for 2 hrs at room temperature after which the FITC-Tax-1 10-mer (50nM) was added and further incubated for 1 hr. The plates were read out in FlexStation 3 (Molecular devices) at 23 °C, using 485 nm excitation and 520 nm emission. For the control FP experiments (Fig. S4), dilution series of GST-hDLG1 PDZ proteins (direct binding), and the dilution series of unlabelled Tax-1 10-mer were assayed as done with the small molecule (competitive binding).

### 2.10. Bio-layer interferometry (BLI)

All experiments were performed in 20 mM sodium phosphate pH 6.0, 50 mM NaCl, 2 mM DTT, and 0.03% DDM in 96 well-plates on the BLI Octet instrument (ForteBio, Inc., USA) at 27 °C. The streptavidin Octet SA biosensors (Sartorius) were loaded with biotinylated Tax-1 10-mer peptide (Biotin-Ahx-SEKHFRETEV-OH) (100 nM solution) for 150-170 s, followed by buffer equilibration for 60 s. An association step of 120 s was sufficient to reach the binding equilibrium as evidenced by flat plateaus of the BLI response curves. Finally, protein dissociation (120 s) and sensor regeneration (5 repeats of 5 s 0.5 M H_3_PO_4_ soak and 5 s buffer wash) completed the measurement cycle. A double referencing with a protein-free sample and an unmodified sensor in the working buffer was performed throughout. For the small molecule screening, the samples contained 5 μM hDLG1 PDZ1 and 100 μM compounds (added from 20 mM stock in DMSO); corresponding protein-free, compound-only reference samples (used for subsequent background signal subtraction) acquired throughout; and a fresh set of biosensors used for each screened compound. For the control BLI experiments (Fig. S5), dilution series of hDLG1 PDZ proteins were assayed in direct binding setup, and dilution series of Tax-1 PBM 4-mer peptide at the constant protein concentration (5 μM) were assayed in competitive binding experiments.

### 2.11. Test for inhibition of cell-to-cell HTLV-1 transmission

Virus-producing cells (MT2) were co-cultured with Jurkat cells containing the gene encoding luciferase under the control of the HTLV-1 LTR5’ viral promoter. The cells were grown for 48 hours in the presence or absence of varying amounts of **3**. Luciferase activation assay was performed as described above.

## 3. Results

### 3.1. The structural basis of Tax-1/ hDLG1 PDZ interactions

Tax-1 is a homodimeric protein bearing a short PDZ-binding motif (PBM) at the C-terminus (Fig. 1A). In turn, hDLG1 harbours three PDZ domains: a closely-spaced PDZ1-PDZ2 tandem and a distant PDZ3 domain (Fig. 1A), all of which can bind full-length Tax-1 (Fig. 1B,C). In the context of the full-length proteins, the interaction is believed to proceed via the hDLG1 PDZ1+2 tandem engaging a pair of PBMs in the Tax-1 homodimer, as seen in other complexes of PDZ tandem-bearing proteins with their dimeric partners (Grembecka et al., 2006). While Tax-1-hDLG1 binding was previously biochemically and functionally validated (Aoyagi et al., 2010; Hirata et al., 2004; Marziali et al., 2017; Suzuki et al., 1999), little is known about its molecular determinants, which is essential for structure-based drug discovery.

**Fig. 1.**
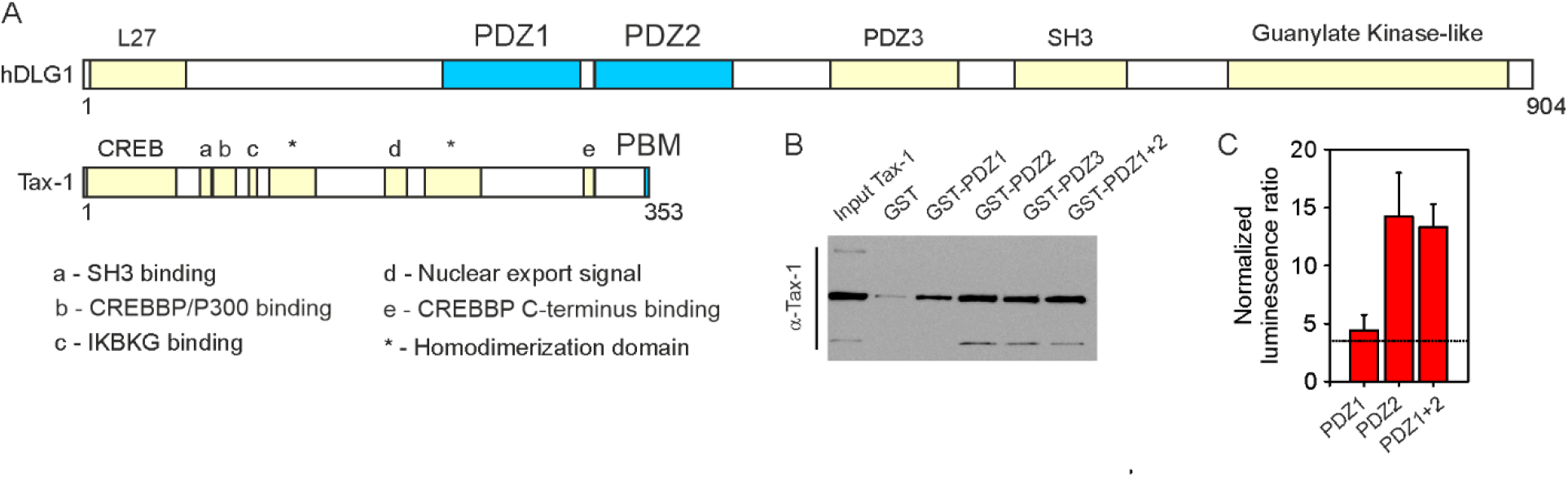
Tax-1-hDLG1 PDZ interactions. **(A)** Domain organization of hDLG1 and Tax-1 proteins. **(B)** Pull-down assays with GST-fused hDLG1 PDZ domains as bait and α-Tax-1 used for detection. **(C)** Mammalian nanoluciferase 2-hybrid (mN2H) assays with N1-fused Tax-1 and N2-fused hDLG1 PDZ domains. The horizontal line denotes the threshold NLR value of 3.5.

To obtain molecular-level details of the Tax-1-hDLG1 interaction, we employed solution NMR spectroscopy. First, we studied binding of the last 10 C-terminal amino acids of Tax-1 (Tax-1 10-mer, H-SEKHFRETEV-OH) to the tandem PDZ1+2 domains of hDLG1. Monitored in a series of [^1^H,^15^N] heteronuclear single-quantum correlation (HSQC) experiments, binding of the natural-abundance, NMR-silent Tax-1 10-mer to the ^15^N labelled, NMR-visible hDLG1 PDZ1+2 leads to large spectral changes (Fig. 2A-B). For both PDZ domains, the strongest effects were observed for the residues in and around the canonical peptide binding site, formed by the β1/β2 loop, the β2 strand and the α2 helix (Fig. 2B). In particular, the largest binding shifts are detected for the PBM residues G235 and F236 of PDZ1 and G330 and F331 of PDZ2 (Fig. 2A-B). The resulting chemical shift perturbation maps show the contiguous protein surfaces of both PDZ1 and PDZ2 targeted by the Tax-1 10-mer (Fig. 2C-D). Moreover, the binding effects of the Tax-1 10-mer to individual PDZ1 (Fig. 2E-G) or PDZ2 (Fig. 2I-K) domains are similar to those observed for the PDZ1+2 tandem (Fig. 2A-D), confirming that separate hDLG1 PDZ domains can function as independent modules. The interaction of Tax-1 10-mer with PDZ2 is stronger than that with PDZ1 (K_D_ of 0.9 μM and 2.7 μM, respectively; Fig. 2H, L), which agrees with GST-pulldown and N2H assays (Fig. 1B,C).

**Fig. 2.**
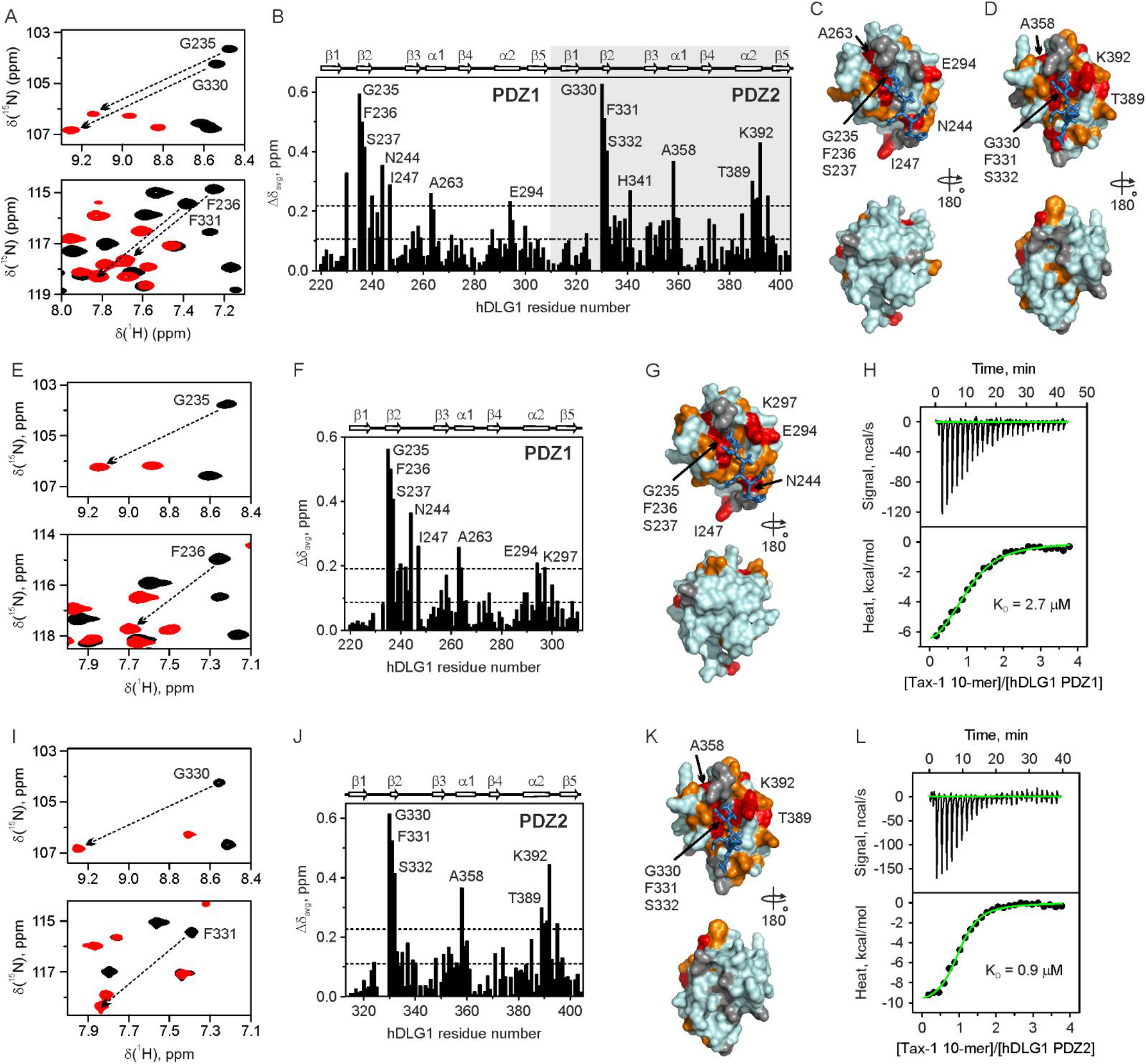
Interaction of Tax-1 10-mer with hDLG1 PDZ domains. NMR-observed peptide binding to the ^15^N hDLG1 **(A-D)** PDZ1+2 tandem or individual **(E-G)** PDZ1 and **(I-K)** PDZ2 domains. **(A,E,I)** Overlay of ^1^H,^15^N HSQC spectral regions in the absence and presence of Tax-1 10-mer peptide (black and red, respectively). **(B,F,J)** Average amide binding shifts (Δδ_avg_) at saturating amounts of Tax-1 10-mer. The horizontal lines show the average Δδ_avg_ and the average (avg) plus one standard deviation (stdev). The secondary structure of the hDLG1 PDZ1 and PDZ2 domains is shown above the plot. **(C,D,G,K)** Chemical shift mapping of the Tax-1 10-mer binding. The molecular surfaces of hDLG1 **(C,G)** PDZ1 and **(D,K)** PDZ2 domains are coloured by the Δδ_avg_ values (orange: Δδ_avg_ > avg; red: Δδ_avg_ > avg + stdev). Prolines and residues with unassigned backbone amide resonances are in grey. The modelled C-terminal part of the Tax-1 peptide, bound to the canonical PDZ site, is shown in sticks. **(H,L)** Binding of the Tax-1 10-mer observed by isothermal titration calorimetry (ITC). The top and bottom panels show, respectively, the raw data after the baseline correction and the integrated data corrected for the heat of Tax-1 10-mer dilution. The solid line in the bottom panel shows the best fit to a binding model with **(H)** the stoichiometry n = 1.03 ± 0.03, equilibrium dissociation constant K_D_ = 2.7 ± 0.3 μM, and the binding enthalpy ΔH = −8.0 ± 0.3 kcal/mol and **(L)** n = 0.99 ± 0.02, K_D_ = 0.9 ± 0.1 μM, and ΔH = - 10.4 ± 0.2 kcal/mol.

To narrow down further the Tax-1 binding requirements, we performed NMR experiments with the peptides corresponding to the last four and first six amino acids of the Tax-1 10-mer. The C-terminal PBM 4-mer (H-ETEV-OH) interacts with the ^15^N hDLG1 PDZ1+2 tandem in the same way as the Tax-1 10-mer (Fig. S1A-D), while no binding of the N-terminal 6-mer peptide (SEKHFR) could be observed (Fig. S1E,F). These findings demonstrate that the four C-terminal residues of Tax-1, bearing the X-S/T-X-V binding motif, are necessary and sufficient for the interaction with the hDLG1 PDZ1 and PDZ2 domains. Furthermore, this interaction requires a free C-terminal carboxyl group as illustrated by smaller chemical shift perturbations and a 360-fold decrease of the binding affinity in NMR experiments with a chemically modified, C-terminally amidated Tax-1 peptide H-ETEV-NH_2_ (Fig. S2).

### 3.2. Crystal structures of hDLG1 PDZ1 and PDZ2 with Tax-1 ETEV peptide

Guided by the above findings, we crystallized complexes of hDLG1 PDZ1 and PDZ2 domains with the ETEV tetramer, the minimal Tax-1 PBM fragment. In both X-ray structures, the peptide sits in a groove formed by the β2-strand, α2-helix, and β1-β2 loop (Fig. 3, left). The latter contains a conserved GLGF motif, which is essential for intermolecular interactions with the C-terminal carboxylate of the ETEV tetrapeptide (Fig. 3, right). In particular, the hydrogen bonding between the terminal carboxyl oxygens of the ETEV peptide with the backbone amide protons in the PDZ GLGF loop stabilizes V353 and orients its side chain inward the binding groove. Thus, both hDLG1 PDZ1 and PDZ2 domains present a hydrophobic binding pocket to accommodate the C-terminal valine of Tax-1 (Fig. S3). In both PDZ-peptide structures, the side chain hydroxyl of T351 forms a hydrogen bond with the imidazole group of histidine in the first position of the α2 helix – a recognition mode typically observed in the class I PDZ domains (Christensen et al., 2019). Rotation of the E350 side chain causes a difference in the interactions of ETEV with the two PDZ domains: while in the PDZ1 complex E350 interacts with threonine in the β3 strand (Fig. 3A), in the PDZ2-ETEV complex it engages a serine in the β2 strand (Fig. 3B). Finally, in both structures, we observe a backbone-backbone interaction between the antiparallel β2-sheet of the PDZ domains and the ETEV peptide Fig. 3, right, a recurrent mechanism in PDZ-peptide binding known as β-augmentation (Heuck et al., 2011).Overall, the hDLG1 PDZ-PBM interactions seen in the X-ray structures fully account for the binding effects observed by NMR spectroscopy (Fig. 2), suggesting that the PDZ-PBM binding mode in the crystal is preserved in solution.

**Fig. 3.**
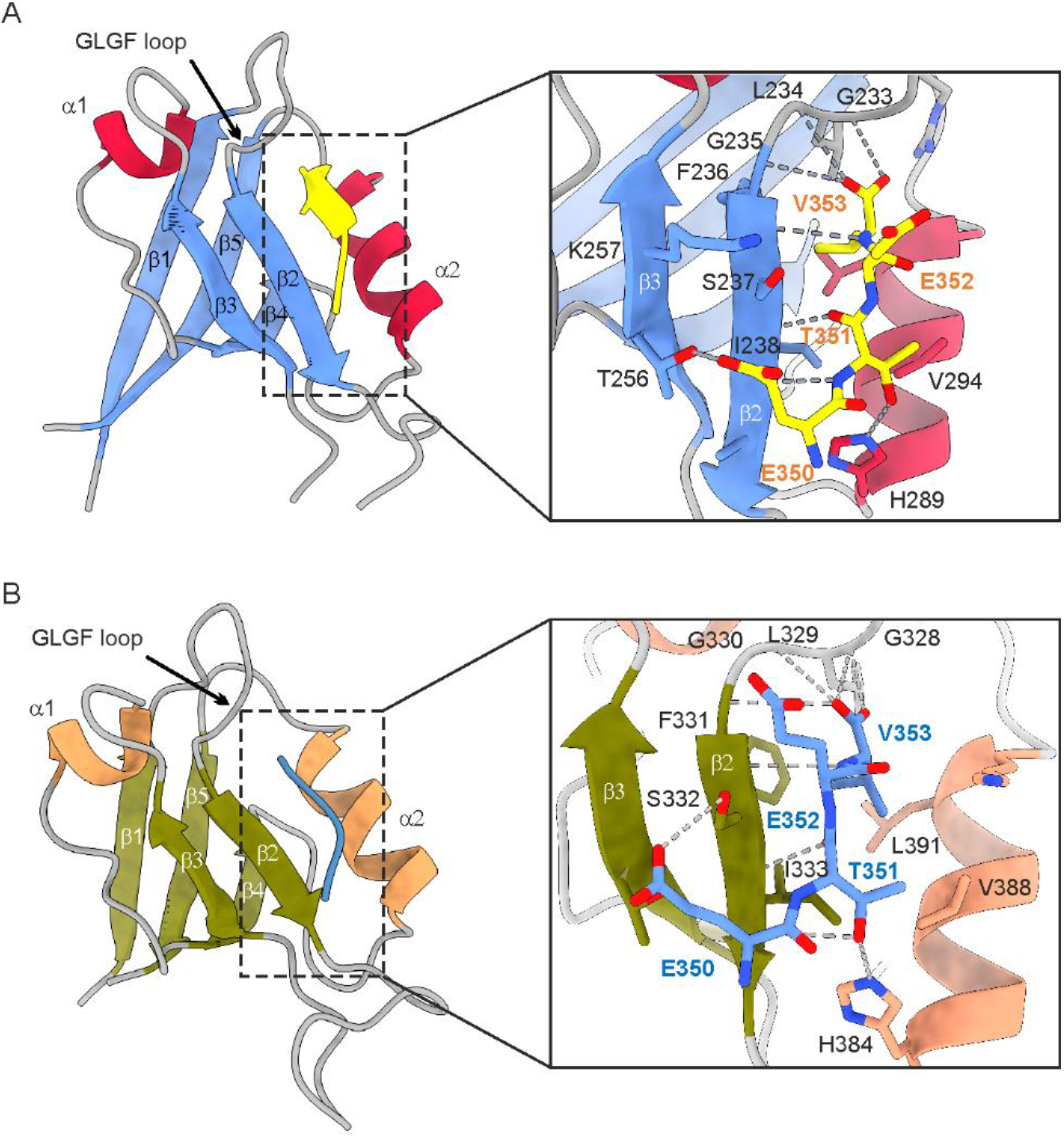
X-ray structures of hDLG1 PDZ1 and PDZ2 domains bound to the Tax-1 PBM 4-mer ETEV peptide. Overall structure of **(A)** PDZ1 and **(B)** PDZ2 complexes in ribbon representation, colored according to the secondary structure. The peptide binding site is delimited by the β2 strand, the α2 helix, and the GLGF loop. (Inset) Close-up view of the ETEV peptide (in stick representation) bound to the PDZ domains. Important residues are labelled and shown in sticks; intermolecular hydrogen bonds are represented by dashed lines.

### 3.3. Small molecule inhibition of Tax-1-hDLG1 interaction

In order to identify potential inhibitors of the Tax-1-hDLG1 interaction, we performed an ultra-large virtual screening of small molecule libraries using crystal structures of hDLG1 PDZ-peptide complexes as an input. In particular, we screened over 10 million commercially-available compounds from the ZINC15 library using VirtualFlow platform (Gorgulla et al., 2020) and selected 212 candidate molecules with high docking scores for further experimental validation. First, these were validated *in vitro* using FP assays, which follow dislocation of the fluorescently-labelled Tax-1 10-mer peptide from pre-formed complexes with hDLG1 PDZ domains. Competitive binding of a small molecule displaces the bound reporter peptide, resulting in FP signal. Out of the 212 tested compounds, 19 showed a clear FP signal decrease of more than 20% (Fig. 4A). These were assayed further by BLI, which monitors interaction of hDLG1 PDZ with Tax-1 10-mer attached to the optical sensor, with the competing small molecules interfering with the binding and, thus, decreasing the BLI response. The FP and BLI assays were fully validated in control experiments that followed both direct and competitive binding of Tax-1 derived peptides to the hDLG1 PDZ domains (Figs. S4 and S5). Non-specific interactions of small molecules with the BLI sensor surface generated a strong background signal, the substraction of which (obtained from the corresponding protein-free samples) resulted in large measurement errors (Fig. 4A). Nonetheless, we could clearly identify several compounds that displayed over 50% decrease in BLI response, a result which was in full agreement with the FP assays (Fig. 4A).

**Fig. 4.**
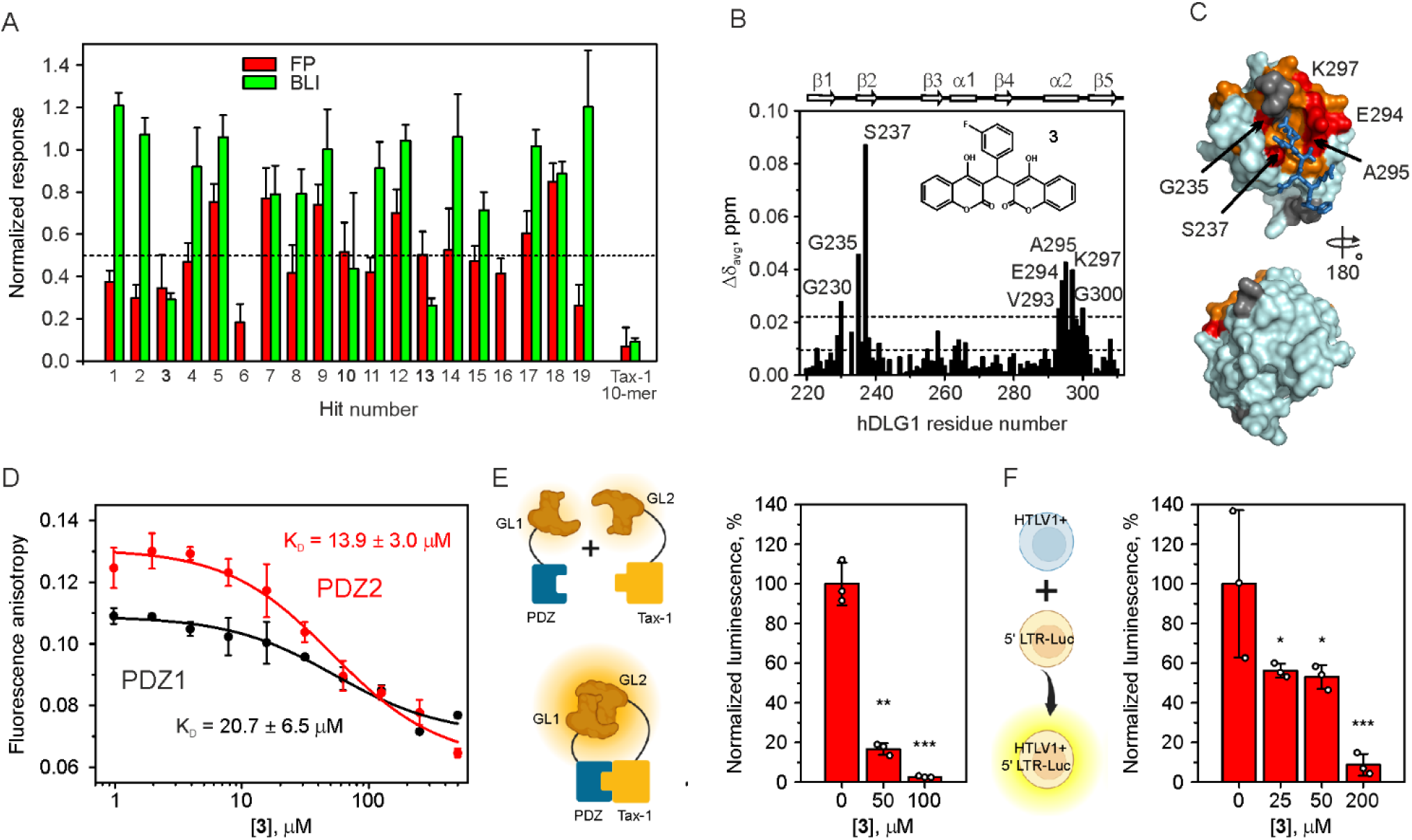
Screening and validation of small molecule inhibitors of Tax-1-hDLG1 interaction. **(A)** Validation of virtual screening hits for hDLG1 PDZ1 binding by FP and BLI. The data for Tax-1 10-mer are included as a positive control. The horizontal line denotes a 50% signal decrease. The small molecules below this threshold are in bold. **(B-C)** NMR-observed binding of **3** to ^15^N hDLG1 PDZ1. **(B)** Δδ_avg_ with 4 molar equivalents of **3**. The horizontal lines indicate the avg and avg + stdev. The secondary structure of the hDLG1 PDZ1 domain is shown above the plot. **(C)** Chemical shift mapping of the **3** binding. The molecular surface of hDLG1 PDZ1 is colored by the Δδ_avg_ values (orange: Δδ_avg_ > avg; red: Δδ_avg_ > avg + stdev). Prolines and residues with unassigned backbone amide resonances are colored grey. The *C*-terminal part of the Tax-1 peptide is shown in sticks as a reference. **(D)** FP competition experiments showing the displacement of the PDZ-bound fluorescently labelled Tax-1 10-mer peptide by **3**. The K_D_ values for **3** binding to PDZ domains are indicated in the plot. **(E)** Inhibition of the Tax-1-hDLG1 interaction by **3** in mammalian cells followed in GPCA assay. **(F)** Inhibition of HTLV-1 virus transmission by **3** in luciferase-based viral transmission assay. The datapoints in **(A, D-F)** represent the mean (n=3); the errors are standard deviations from the mean. Statistical significance in **(E)** and **(F)** was determined with a two-tailed t test (*p <0.05, **p <0.01, and ***p <0.001).

One of these small molecules (**3**) showed consistent results in follow-up biophysical and cellular experiments. In particular, NMR spectroscopy revealed that its binding effects on hDLG1 PDZ1 domain are very similar to those of the cognate Tax-1 10-mer peptide (Fig. 4B-C, cf. Fig. 2F-G). Just as Tax-1 PBM, this compound engages the PDZ peptide-binding groove, confirming it as a competitive inhibitor of the Tax-1-hDLG1 PDZ interaction. Furthermore, as observed in the FP competition experiments, this small molecule binds to both hDLG1 PDZ1 and PDZ2 domains, with affinity constants (K_D_) in the 14-21 μM range (Fig. 4D). Finally, **3** exhibits a clear, concentration-depended inhibition of the full-length Tax-1-hDLG1 PDZ interaction in mammalian cells (Fig. 4E) and inhibits HTLV-1 virus transmission in a luciferase-based cell-to-cell transmission assay (Fig. 4F).

## 4. Discussion

Despite the discovery of viruses in the early 1900s, and the demonstration of their pathogenicity in humans in early 1950s, the arsenal of antiviral drugs remains dangerously small, with only ~90 molecules available today being biased towards viral enzymes (De Clercq and Li, 2016). Furthermore, 95% of approved drugs target only 6 types of viruses (HIV-1, HCV, HBV, HSV, HCMV, and Influenza viruses), with HIV-1 covering the majority of the available drugs. As evidenced by the coronavirus disease 19 (COVID19) pandemic, it is of utmost importance to anticipate emerging and re-emerging viral infections by identifying potential, broad-spectrum antiviral drugs. However, this task is rendered difficult by many challenges, among which are: (i) the druggability of viral products, (ii) the antiviral drug safety, and (iii) a desirable high potency to limit the selection of resistant mutations. High-throughput mapping of host-virus PPIs and massively parallel genomic sequencing have accelerated the pace of discovery of key host targets, which are shared by different categories of pathogenic viruses, including RNA and DNA, or acute and persistent viruses (Rozenblatt-Rosen et al., 2012; Tang et al., 2013; Watanabe et al., 2014). These host factors provide novel opportunities to prioritize antiviral drugs based on common host targets. In this work, our goal was to demonstrate that one of such host determinants, the PDZ-domain containing proteins, could provide insights into how to identify small molecule inhibitors of viral-host PPIs.

The methodology used here defines a pipeline that could be applied to any pathogenic parasite of interest, without prior knowledge on the druggability of its encoded gene products. First, we chose HTLV-1 Tax (Tax-1), a highly connected viral oncoprotein(Vandermeulen et al., 2021). Second, we determined the structural basis of Tax-1 C-terminal motif binding to canonical sites of hDLG1 PDZ domains. Finally, we demonstrated that a small molecule targeting Tax-1-hDLG1 interactions could inhibit Tax-1 functions in cellular models.

hDLG1 has also been shown to interact with HPV E6, Ad9 E4orf1, IAV NS1, HTLV-1 Env, HIV-1 Env and HCV core proteins, but with different biological outcomes, including perturbations of cell polarity, signalling, or apoptosis (Thomas and Banks, 2018). Interestingly, the conserved C-terminal PBM motifs of these viral proteins highly correlate with viral pathogenicity, independently of their substantial differences in genome content (RNA and DNA viruses) or mode of infection (acute and persistent families of viruses). hDLG1 is thus a “pan-viral interactor” and may constitute an ideal drug target for the discovery of broad-spectrum antiviral inhibitors.

Structural insights are essential for design of small molecule inhibitors of interactions between viral proteins and cellular protein modules. The X-ray structures of hDLG1 PDZ1 and PDZ2 complexes with Tax-1 PBM peptide reported in this work show that their C-terminal carboxyl groups make hydrogen bonds with the conserved GLGF loop residues (G233-F236 and G328-F331 in hDLG1 PDZ1 and PDZ2, respectively), in agreement with other structural studies using a Tax-1-unrelated X-S/T-X-V motif (Zhang et al., 2011). Comparison across different PDZ-peptide systems shows that hDLG1 residues G235, F236, G330, and F331 act as binding hot spots, which should be amenable to traditional drug discovery efforts aiming at designing potent small molecule binders. Interestingly, an X-ray structure of the hDLG1 PDZ2 with the C-terminal Tax-1 10-mer peptide has been published recently (Cousido-Siah et al., 2022), showing essentially the same peptide binding mode as the one reported here. The main difference is the orientation of the E350 side chain, which interacts with the threonine in the β3 strand in that work, as opposed to the serine in the β2 strand seen here. In addition, the published structure with a longer Tax-1 peptide features an interaction between the backbone of R349 and the N339 side chain, which likely further stabilizes the peptide binding.

As a proof-of-concept, we demonstrated a clear correlation between this PPI inhibition and impairment of Tax functions, including its transformation ability and HTLV-1 cell-to-cell transmission. The combination of our experimental strategies should allow the development of anti-Tax-1 compounds for future pre-clinical and clinical studies of drug candidates in view of future treatments of HTLV-1-induced diseases.

## Author contributions

Conceptualization, S.M., J-C.T., I.VM. and A.V.; Methodology, S.M., Y.B, I.VM., A.V, C.G. and J-C.T; Investigation, S.M, Y.B, I.VM, C.G.,C.M., J.O.,J.B., T.N., A.S., A.V. and J-C.T.; Formal analysis, S.M., J-C.T., I.VM. and A.V. Resources, H.A., A.V. S.B., F.D, H.R, K.S.A, and J-C.T.; Writing - Original Draft, S.M., I.VM, A.V. and J-C.T.; Writing, - Review and Editing, J.O., A.V., K.S.A and J-C.T.; Supervision, J-C.T., A.V., H.R, S.B, K.S.A; Funding Acquisition, S.B, H.A, K.S.A and J-C.T.

## Data and materials availability

All data and materials used in the analyses are described in this manuscript. Plasmids and vectors are available via materials transfer agreements (MTAs).

## Supplementary

**Fig. S1.**
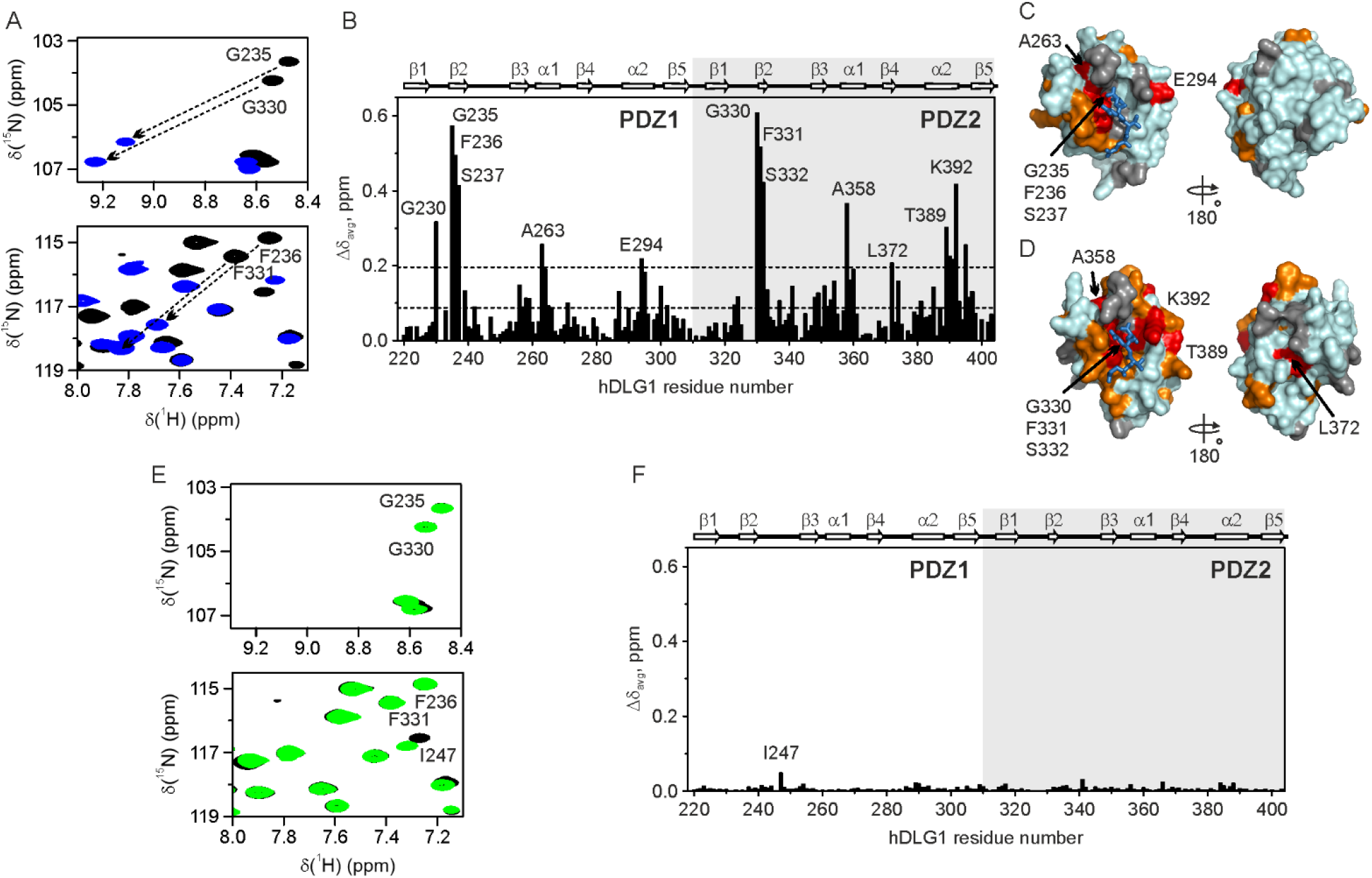
Control hDLG1 PDZ NMR experiments with Tax-1 PBM 4-mer and N-terminal 6-mer (6-mer-N) peptides. **(A)** Overlay of ^1^H,^15^N HSQC spectral regions of the free and PBM 4-mer-bound ^15^N hDLG1 PDZ1+2 tandem (in black and blue, respectively), with the chemical shift perturbations for the selected residues indicated by arrows. **(B)** Δδ_avg_ of the hDLG1 PDZ1+2 at saturating amounts of PBM 4-mer. **(C,D)** Chemical shift mapping of the PBM 4-mer binding. The molecular surfaces of hDLG1 **(C)** PDZ1 and **(D)** PDZ2 domains are colored as in Fig. 2C,D. The modeled PBM peptide, bound to the canonical PDZ site, is shown in sticks. **(E)** The spectra of the ^15^N hDLG1 PDZ1+2 tandem in the absence and presence of 6-mer-N (black and green, respectively). **(F)** Δδ_avg_ of the hDLG1 PDZ1+2 in the presence of 6 molar equivalents of 6-mer-N.The labels identify the sole residue with a significant binding shift.

**Fig. S2.**
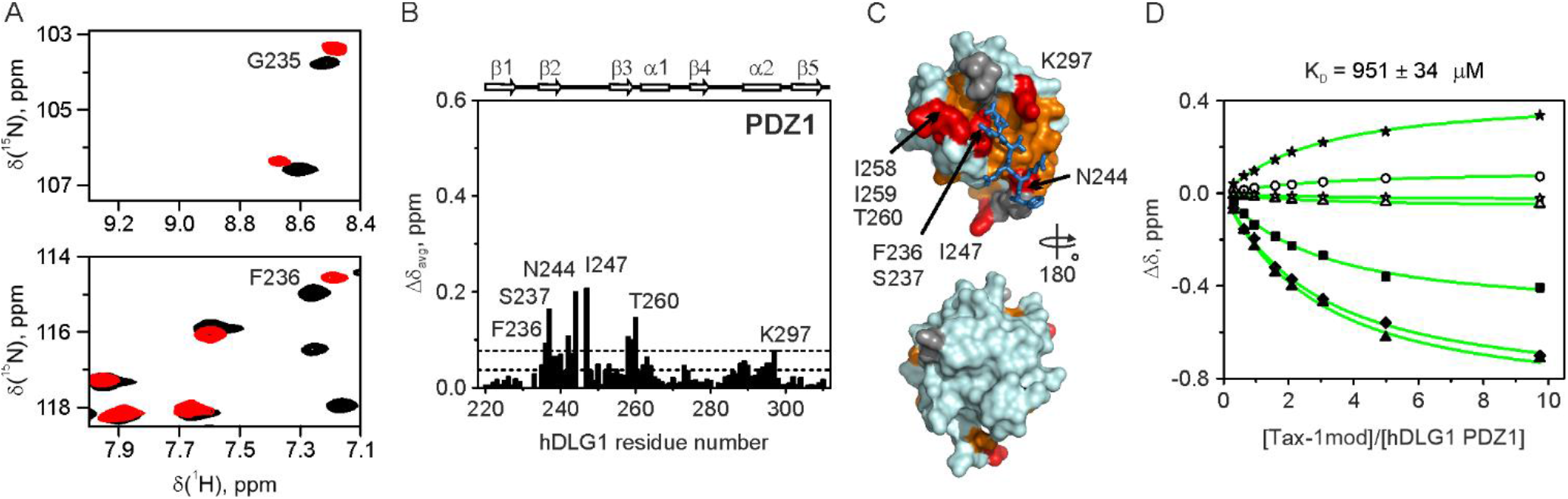
Control NMR experiments with the C-terminally amidated Tax-1 10-mer (Tax-1mod). **(A)** Overlay of selected ^1^H,^15^N HSQC spectral regions of ^15^N hDLG1 PDZ1 in the absence and presence of Tax-1mod peptide (black and red, respectively). The labels identify the backbone amide resonances affected by the Tax-1 10-mer binding (cf. Fig. 2). **(B)** Δδ_avg_ of hDLG1 PDZ1 at saturating amounts of Tax-1mod. The horizontal lines show the average Δδ_avg_ and the avg + stdev. Residues with large Δδ_avg_ are labelled. The secondary structure of hDLG1 PDZ1 is shown above the plot. **(C)** Chemical shift mapping of the Tax-1mod binding. The molecular surface of hDLG1 PDZ1 is colored by the Δδ_avg_ values (orange: Δδ_avg_ > avg; red: Δδ_avg_ > avg + stdev). Prolines and residues with unassigned backbone amide resonances are in grey. The modeled C-terminal part of the Tax-1 PBM, bound to the canonical PDZ site, is shown in sticks. **(D)** NMR chemical shift titration of ^15^N hDLG1 PDZ1 with the Tax-1mod peptide. Open and filled symbols refer to the chemical shift perturbations (Δδ) of the backbone H and N atoms, respectively, of F236 (squares), G240 (circles), I258 (triangles), I259 (diamonds), and L296 (stars). The NMR titration curves were analyzed with a two-parameter nonlinear least squares fit with the shared K_D_ using a one-site binding model corrected for the dilution effect(Kannt et al., 1996). The solid lines show the best fits with the K_D_ value of 951 ± 34 μM.

**Fig. S3.**
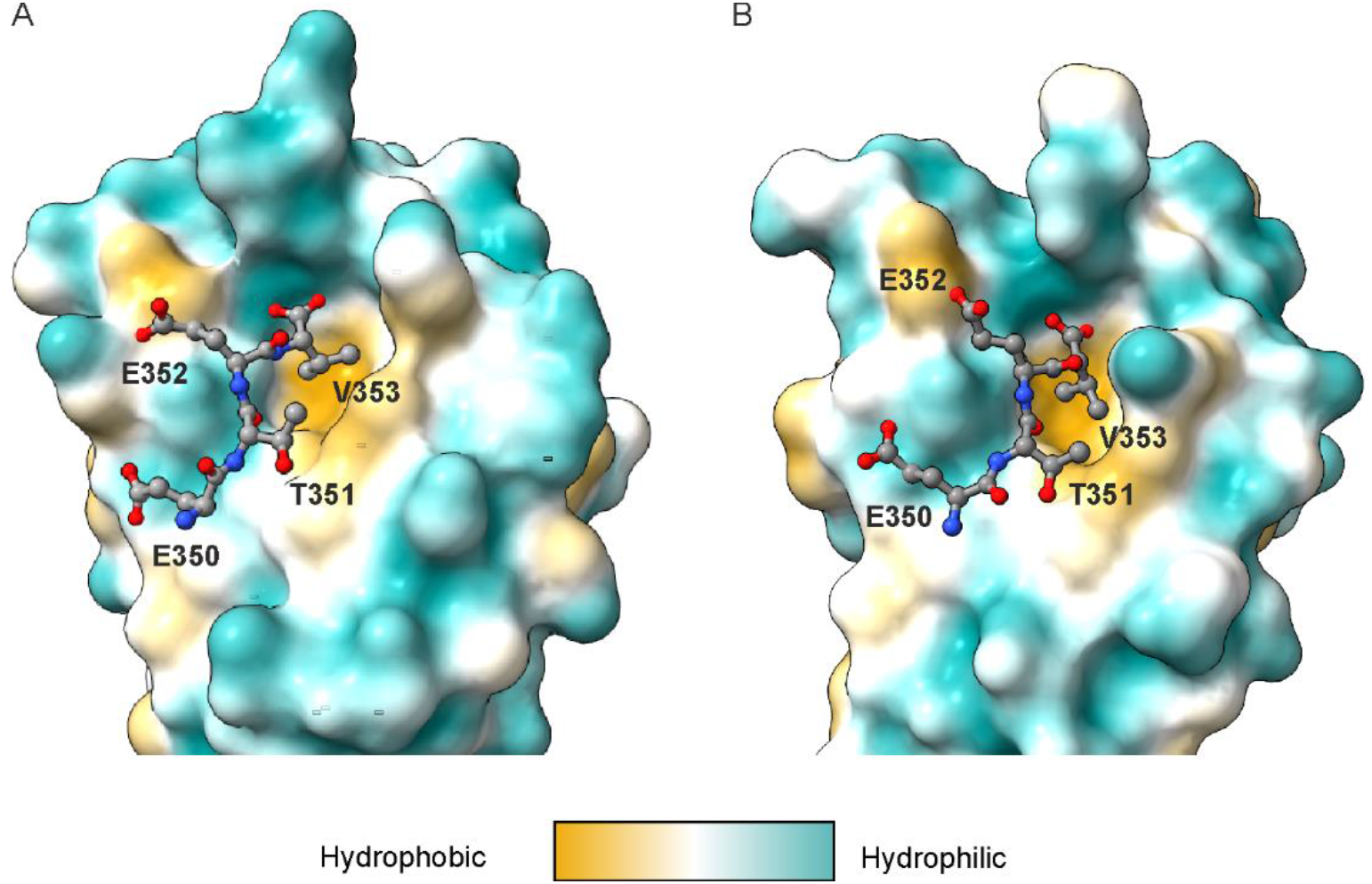
Hydrophobicity of the binding groove in hDLG1 **(A)** PDZ1 and **(B)** PDZ2 complexes with the Tax-1 PBM ETEV tetrapeptide.

**Fig. S4.**
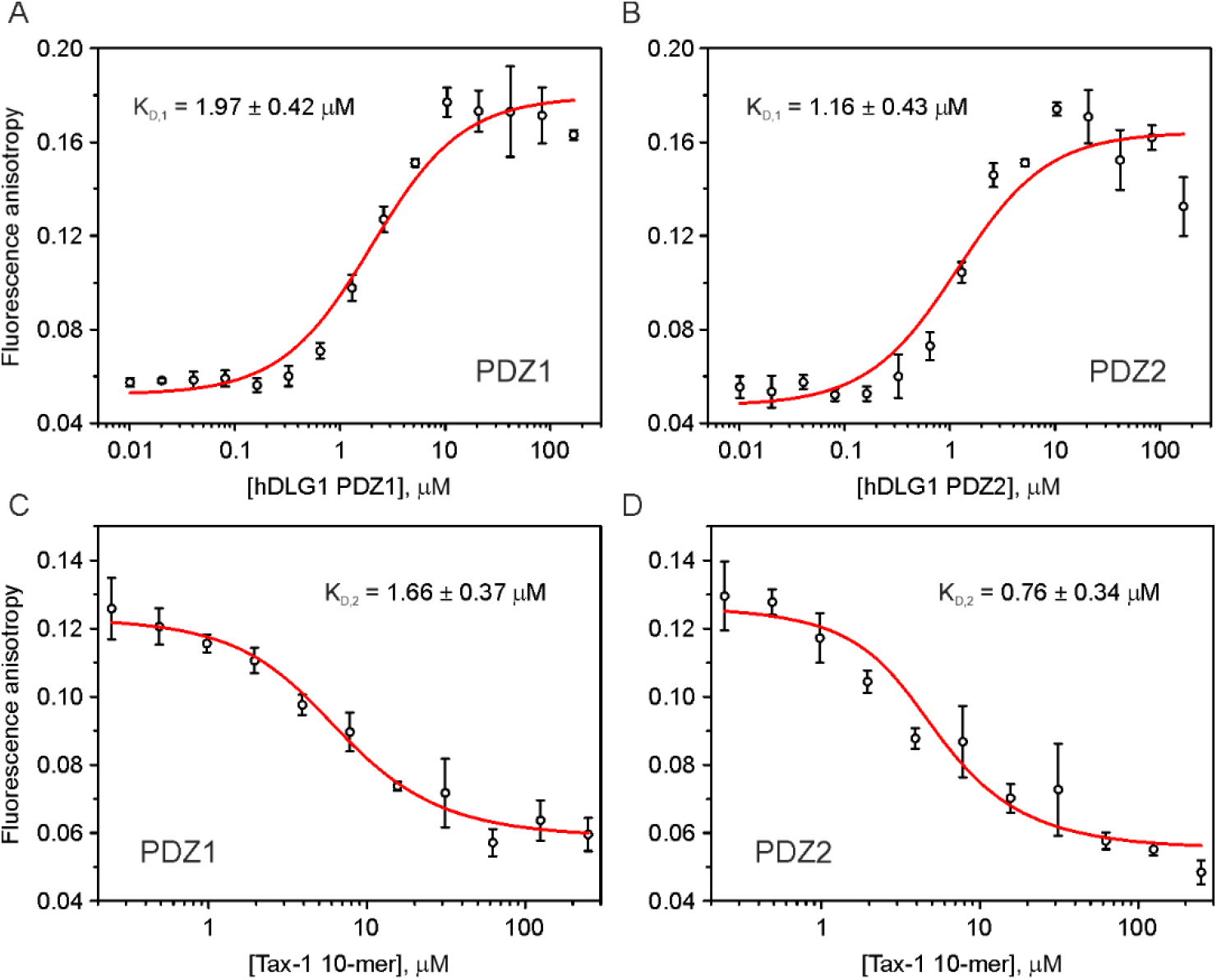
Control FP experiments. **(A,B)** Direct binding of the fluorescently-labelled Tax-1 peptide (FITC-Tax-1 10-mer) to hDLG1 **(A)** PDZ1 and **(B)** PDZ2 domains. **(C,D)** Competition experiments with unmodified Tax-1 10-mer peptide titrated into the preformed complex of FITC-Tax-1 10-mer with hDLG1 **(C)** PDZ1 and **(D)** PDZ2 domains. The red curves are the best fits to direct and competitive FP binding models(Roehrl et al., 2004), with K_D_ values indicated in the plots. The protein concentration in competition experiments was 3 μM; the concentration of the reporter FITC-Tax-1 10-mer was 50 nM; throughout. All experiments were performed in triplicates at 23 °C in 20 mM sodium phosphate pH 6.0, 50 mM NaCl, 2 mM DTT. The errors are standard deviations from the mean.

**Fig. S5.**
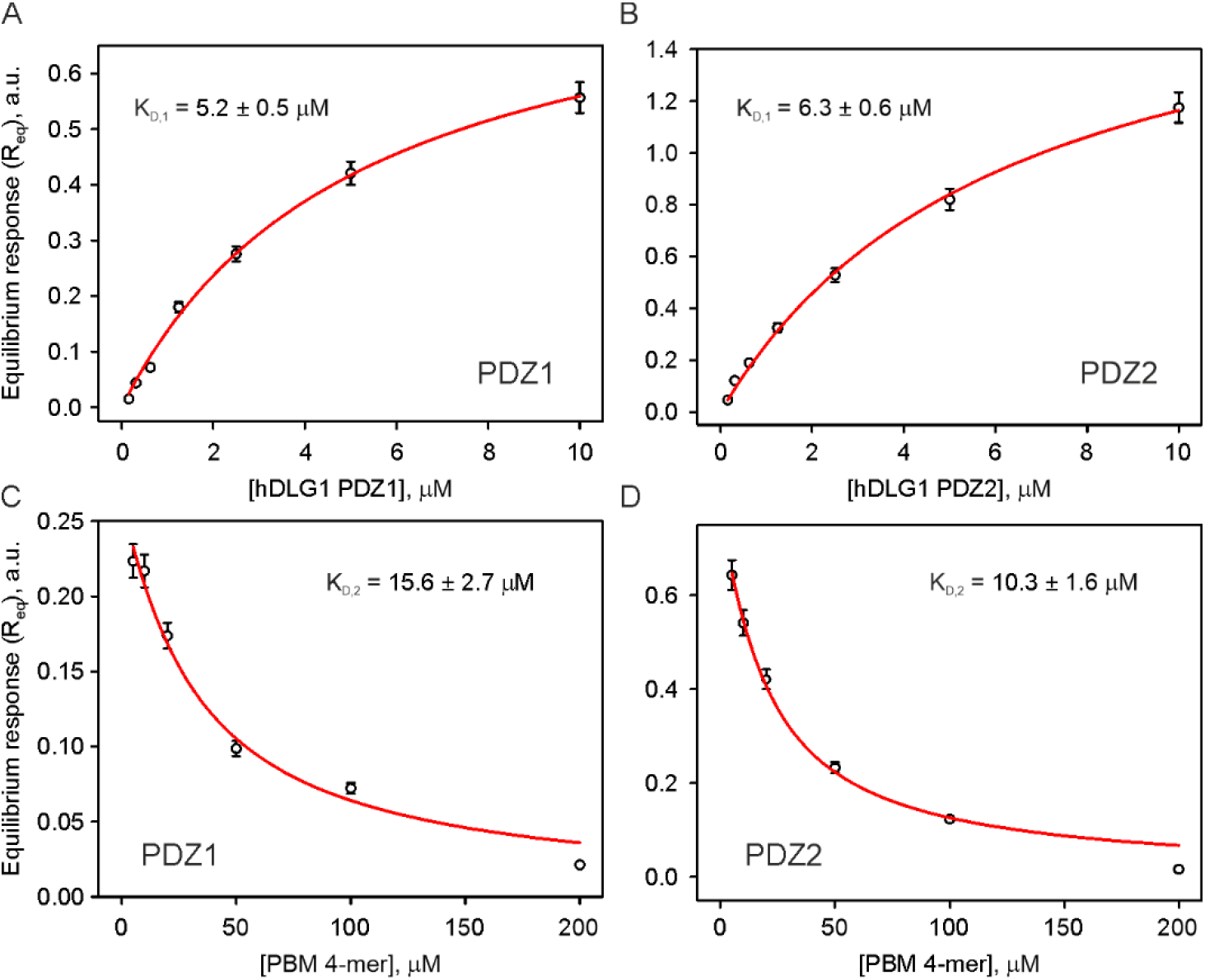
Control BLI experiments. **(A,B)** Direct binding of hDLG1 **(A)** PDZ1 and **(B)** PDZ2 domains to the BLI sensor modified with Tax-1 10-mer peptide. **(C,D)** Competition experiments with PBM 4-mer ETEV peptide binding to hDLG1 **(C)** PDZ1 and **(D)** PDZ2 domains. The red curves are the best fits to direct and competitive binding models (equations 1 and 5, respectively, in Supplementary Text 2), with K_D_ values indicated in the plots. All experiments were performed at 27 °C in 20 mM sodium phosphate pH 6.0, 50 mM NaCl, 2 mM DTT, and 0.03% DDM. The protein concentration in competition experiments was 5 μM. The error bars denote standard measurement errors of 5%.

**Table S1.**
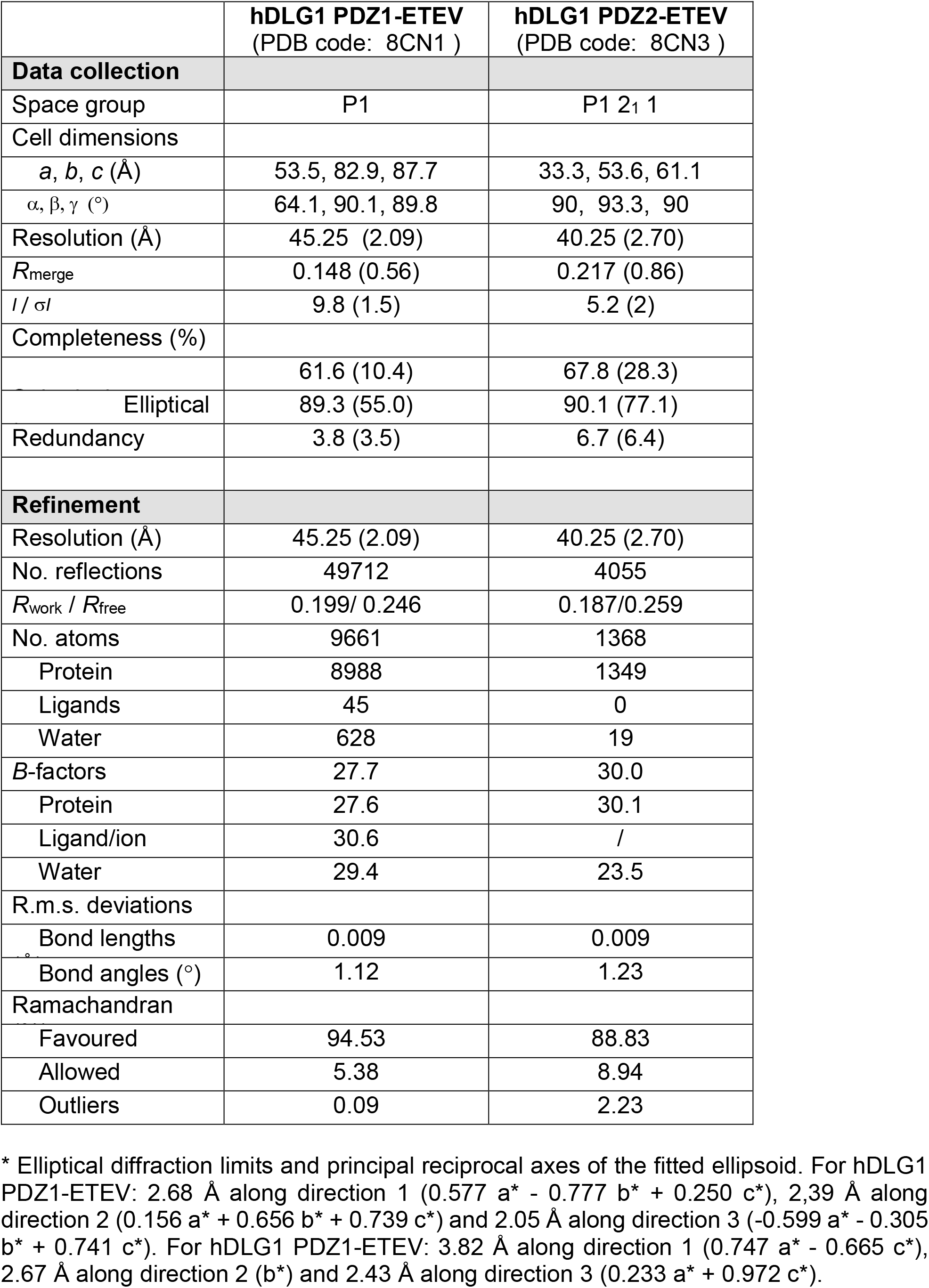
X-ray crystallography data collection and refinement statistics.

|  | **hDLG1 PDZ1-EDEV**<br>(PDB code: 8CN1 ) | **hDLG1 PDZ2-EDEV**<br>(PDB code: 8CN3 ) |
| --- | --- | --- |
| **Data collection** |  |  |
| Space group | P1 | P1 2<sub>1</sub> 1 |
| Cell dimensions |  |  |
| *a*, *b*, *c* (Å) | 53.5, 82.9, 87.7 | 33.3, 53.6, 61.1 |
| $\alpha$, $\beta$, $\gamma$ (°) | 64.1, 90.1, 89.8 | 90, 93.3, 90 |
| Resolution (Å) | 45.25 (2.09) | 40.25 (2.70) |
| *R*<sub>merge</sub> | 0.148 (0.56) | 0.217 (0.86) |
| *I* / σ*I* | 9.8 (1.5) | 5.2 (2) |
| Completeness (%) |  |  |
|  | 61.6 (10.4) | 67.8 (28.3) |
| Elliptical | 89.3 (55.0) | 90.1 (77.1) |
| Redundancy | 3.8 (3.5) | 6.7 (6.4) |
| **Refinement** |  |  |
| Resolution (Å) | 45.25 (2.09) | 40.25 (2.70) |
| No. reflections | 49712 | 4055 |
| *R*<sub>work</sub> / *R*<sub>free</sub> | 0.199/ 0.246 | 0.187/0.259 |
| No. atoms | 9661 | 1368 |
| Protein | 8988 | 1349 |
| Ligands | 45 | 0 |
| Water | 628 | 19 |
| *B*-factors | 27.7 | 30.0 |
| Protein | 27.6 | 30.1 |
| Ligand/ion | 30.6 | / |
| Water | 29.4 | 23.5 |
| R.m.s. deviations |  |  |
| Bond lengths | 0.009 | 0.009 |
| Bond angles (°) | 1.12 | 1.23 |
| Ramachandran |  |  |
| Favoured | 94.53 | 88.83 |
| Allowed | 5.38 | 8.94 |
| Outliers | 0.09 | 2.23 |

\* Elliptical diffraction limits and principal reciprocal axes of the fitted ellipsoid. For hDLG1 PDZ1-EDEV: 2.68 Å along direction 1 (0.577 *a*\* - 0.777 *b*\* + 0.250 *c*\*), 2.39 Å along direction 2 (0.156 *a*\* + 0.656 *b*\* + 0.739 *c*\*) and 2.05 Å along direction 3 (-0.599 *a*\* - 0.305 *b*\* + 0.741 *c*\*). For hDLG1 PDZ2-EDEV: 3.82 Å along direction 1 (0.747 *a*\* - 0.665 *c*\*), 2.67 Å along direction 2 (*b*\*) and 2.43 Å along direction 3 (0.233 *a*\* + 0.972 *c*\*).

### Supplementary Text 1. Synthesis and characterization of biotinylated Tax-1 10-mer peptide

#### Synthesis

Biotin-Tax-1-10-mer peptide (Biotin-Hex-SEKHFRETEV-COOH) was synthesized using Fmoc-based solid phase peptide synthesis (SPPS) on a microwave assisted peptide synthesizer (CEM Liberty Lite). The synthesis was performed on 0.1 mmol scale using preloaded Fmoc-Val-Wang resin. Fmoc deprotection was performed at 90°C for 3 min using a solution of 20% 4-methylpiperidine in DMF during the entire synthesis. Each coupling was done using 5 equivalents of Fmoc protected amino acid, with 0.5 M DIC and 1 M Oxyma as coupling reagents. Then the N-terminus has been functionalized manually through the coupling of 6-Biotinylamino-hexanoic acid (1.5 equivalents) using HBTU/DIPEA in DMF as coupling mixture for 2 h. After completion of the sequence, the resin was washed three times with DCM, followed by the cleavage using a cocktail solution consisting of 90 % TFA, 5 % triisopropylsilane and 5 % distilled water during 4 h. After freeze-drying, crude peptides were obtained and purified using preparative HPLC. More specifically, a Gilson HPLC system, equipped with Gilson 322 pumps and a Supelco Discovery® BIO Wide Pore C18 column (25 cm x 21.2 mm, 10 μm), was used. Crude peptide was dissolved in DMSO and purified using H2O-AcN–0.1% TFA as mobile phase. Finally, fractions were collected, and purity was assessed by analytical RP-HPLC, after which the pure fractions were combined and lyophilized to obtain the final purified peptide as a powder (TFA salt) with a high purity.

HPLC analysis was performed on a Hitachi Chromaster system (Chromaster HPLC 5260 autosampler, Chromaster HPLC 5160 Pump, Chromaster HPLC 5310 column and a Chromaster HPLC 5430 diode array detector). The mobile phase consists of 0.1 % TFA in AcN and 0.1 % TFA in Milli-Q water. The analyzed peptides eluted through a Chromolith® High Resolution C18 end-capped column (50 mm x 4.6 mm, 1.1 μm, 150 Å) using a gradient from 1 % to 100 % of AcN over 6 min at a flow rate of 2.8 mL min^-1^. LC-MS analysis was performed on a Waters 600 HPLC (combined with a Waters 2487 UV detector at 215 nm) connected to a Micromass QTOF-micro system. The mobile phase consists of 0.1 % FA in AcN and 0.1 % FA in Milli-Q water The analyzed peptide eluted through an EC NUCLEODUR C18 endcapped column (150 mm x 2 mm, 5 μm, 300 Å), using a gradient from 3% to 100% AcN over 20 min with a flow rate of 0.3 mL/min.

#### Characterization

Formula: C_70_H_109_N_19_O_22_S

Mw(TFA salt)= 1941 g/mol

HPLC: rt=2.49 min

MS (m/z): [M+H]^+^= 1600.76

yield: 24%

purity > 99%

### Supplementary Text 2. BLI binding models

For direct binding of an analyte in solution (hDLG1 PDZ) to a ligand immobilized on the BLI sensor surface (Tax-1 10-mer peptide), the equilibrium BLI response (R_eq_) is given by Eq. 1:

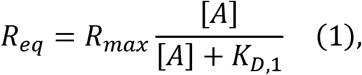

where R_max_ is the maximal BLI response at the ligand saturation, [A] is the analyte concentration, and K_D,1_ is the equilibrium dissociation constant, numerically equal to the analyte concentration at 50% R_max_ (FortéBio). Thus, the direct binding curves in Fig. S5A,B can be fitted with Eq. 1, setting R_eq_ and [A]=[hDLG1 PDZ] as the dependent and independent variables, respectively, and R_max_ and K_D,1_ as the fitted parameters.

For competition experiments, where a competitor (PBM 4-mer) interacts with the analyte (hDLG1 PDZ) in solution, thereby inhibiting its binding to the sensor-immobilized ligand (Tax-1 10-mer peptide), the equation can be derived as follows. For the 1:1 analyte-competitor binding, the dissociation constant (K_D,2_) is given by Eq. 2:

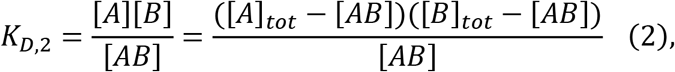

where [A], [B], and [AB] are the equilibrium concentrations of the analyte, competitor, and the analyte-competitor complex, respectively; and [A]_tot_ and [B]_tot_ are the total analyte and competitor concentrations in the sample. Solving the quadratic Eq. 2 gives the expression for the concentration of the analyte bound by the inhibitor, Eq. 3:

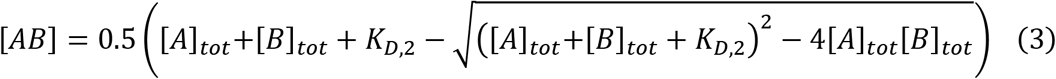

Thus, the concentration of the remaining, free analyte (in solution), available for the interaction with the ligand (on the BLI sensor) can be expressed as:

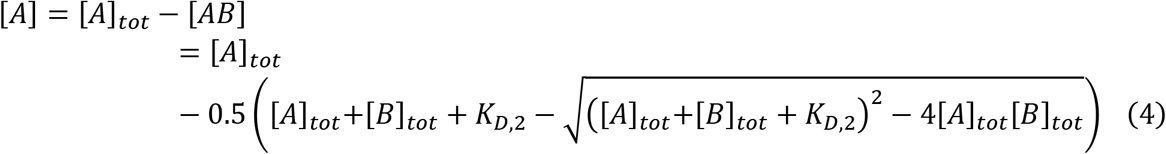

Feeding the expression (4) into Eq. 1 gives the final equation for the BLI competition binding experiment, Eq. 5:

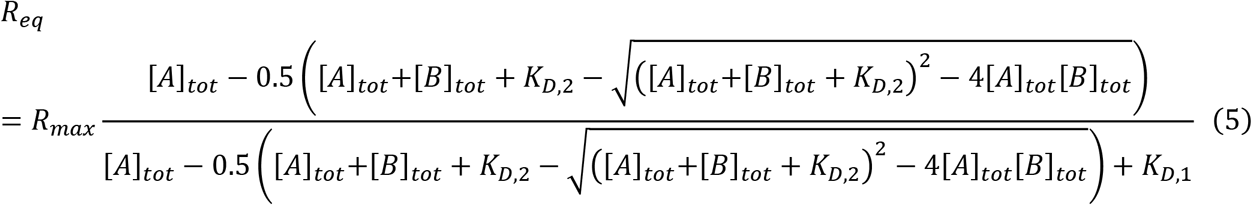

Thus, for the competition experiments in Fig. S5C,D, in which the total analyte concentration [A]_tot_ = [hDLG1 PDZ] is kept constant and that of the inhibitor [B]_tot_ = [PBM 4-mer] is varied (Fig. S5C,D), the binding curves can be fitted with Eq. 5, setting R_eq_ and [B]_tot_ = [PBM 4-mer] as the dependent and independent variables, respectively, and R_max_ and K_D,2_ as the fitted parameters and using the K_D,1_ values determined in the direct BLI binding experiments (Fig. S5A,B).

